# The Impact of *Parabacteroides distasonis* Colonization on Hosts’ Microbiome, Metabolome, Immune Responses, and Diabetes Onset

**DOI:** 10.1101/2024.06.13.598927

**Authors:** Khyati Girdhar, Audrey Randall, Yusuf Dogus Dogru, Clarissa Howard, Alessandro Pezzella, Juan Henao, Umesh K Gautam, Pavla Šašinková, Tomas Hudcovic, Martin Schwarzer, Michael Kiebish, Emrah Altindis

**Affiliations:** Boston College Biology Department, Chestnut Hill, Massachusetts, USA; BPGbio, Inc., Old Connecticut Path, Framingham, MA 01701; Laboratory of Gnotobiology, Institute of Microbiology of the Czech Academy of Sciences, 54922 Novy Hradek, Czech Republic

## Abstract

Type 1 Diabetes (T1D) is a chronic disease caused by autoimmune destruction of insulin-producing pancreatic β-cells. The insulin B-chain 9-23 (insB:9-23) peptide is established as a critical epitope in triggering T1D. In our previous study, we showed that *Parabacteroides distasonis*, a human gut commensal, contains an insB:9-23 mimic in its hprt protein (residues, 4-18). This mimic (hprt4-18) activates insB:9-23 specific T-cells, and colonization of *P. distasonis* in female NOD mice enhanced diabetes onset. Additionally, the presence of hprt:4-18 sequence in the gut microbiome is associated with seropositivity in infants. However, the impact of the colonization on the gut microbiome and intestinal immune cell compositions, gut permeability, cytokine, and serum metabolome profiles were unknown. Here, we addressed this gap using specific pathogen-free (SPF) and germ-free (GF) NOD mouse models. *P. distasonis* colonization had a minimal impact on gut microbiome composition and merely altered 28 ASVs upon colonization. In intraepithelial lymphocytes (IELs) of *P. distasonis* colonized SPF NOD mice, we observed a 1.72-fold reduction in T-helper cells and a 2.3-fold reduction in T-effector cells, along with a 1.85-fold reduction in B-cell populations. Further, *P. distasonis* did not alter serum metabolome and cytokine levels except for a decrease in IL-15. We observed no difference in the gene expression related to gut permeability. Similar to SPF mice, *P. distasonis* colonization in GF NOD mice induced severe insulitis without affecting gut permeability. On the other hand, *P. distasonis* lysate could induce insB:9-23 specific T cells. Altogether, these findings demonstrate that *P. distasonis* does not stimulate a nonspecific inflammatory immune response in the intestines, nor does it cause significant alterations in the gut microbiome, gut permeability, serum metabolome, or cytokine response. However, it does induce insulitis in GF NOD mice and activates insB:9-23 specific T-cells. These findings support our original hypothesis that *P. distasonis* colonization stimulates a specific immune response and enhances T1D onset in NOD mice via molecular mimicry.

## 1. Introduction

Type 1 diabetes (T1D) is an autoimmune disease characterized by the selective destruction of pancreatic β-cells by autoreactive T cells^1^. The incidence of T1D in children is rising annually, with an increase of 3.4% in Europe and 1.4% in the USA^2^ ^3^. Genome-wide association studies have identified over 60 loci that influence the risk of developing T1D^4^; however, genetics alone cannot account for the increasing incidence rates. Various environmental factors, including diet, birth mode, infections, and antibiotics, have also been studied^5–7^. However, the trigger of T1D autoimmunity remains elusive.

Gut microbiota has been increasingly understood over the last 20 years for its role in health and disease. The environmental factors mentioned above can induce functional and compositional changes in the gut microbiota^8^. Several studies have highlighted the continuous crosstalk between the immune system and gut microbes starting immediately after birth^9^. Microbial colonization and exposure to self- and non-antigens shape the host immune system early in life^10^. This coincides with the period when the incidence of T1D is most common. Previous reports have documented changes in gut microbiome composition and diversity in individuals with T1D^11^. The DIABIMMUNE study, a longitudinal examination of the fecal microbiome in HLA and age-matched infants, revealed that there was a higher abundance of pathobionts such as *Ruminococcus gnavus* in seroconverted children compared to the seronegative subjects^12^. In a follow-up study, they found a higher prevalence of *Bifidobacterium* in Russians and LPS-producing *Bacteroides* species in Finns and Estonians. Interestingly *Bacteroides* LPS inhibited innate immune signaling and endotoxin tolerance and *Bacteroides dorei* LPS did not reduce autoimmune diabetes incidence in non-obese diabetic (NOD) mice^13^. Strain-level analysis also revealed significant differences for *Bifidobacterium species*^14^.

The Environmental Determinants of Diabetes in the Young (TEDDY) study, another key longitudinal cohort, used samples from 903 high-risk infants reported, *Parabacteroides* as the only genus significantly associated with the T1D onset^15^. Similarly, the Innovative Approaches to Understanding and Arresting Type 1 Diabetes (INNODIA) study identified *Parabacteroides distasonis* as one of the 30 most abundant species in newly diagnosed individuals^16^. These longitudinal studies offer valuable insights into gut microbiome changes, identifying specific alterations for specific gut commensals, including a significant association between *Parabacteroides* and T1D before disease onset. However, they are descriptive and do not establish causality. *P. distasonis* is a gram-negative, strictly anaerobic gut commensal of humans and other animals. *P. distasonis* was previously reported for its beneficial effects in alleviating inflammatory arthritis^17^, colitis^18^, type 2 diabetes^19^, obesity^20^, non-alcoholic steatohepatitis (NASH)^21^, chronic abdominal pain^22^, and tumorigenesis^23^ ^24^ in different mouse models. *P. distasonis* colonization also increased intestinal barrier integrity and modulated inflammatory markers in A/J mice^25^.

In our previous study^26^, we identified an insB:9-23 mimic in the hypoxanthine phosphoribosyltransferase (hprt) protein of *Parabacteroides distasonis* (hprt4-18). Insulin B chain amino acids 9-23 (<u>insB:9-23</u>)^27^ is one of the most immunodominant T-cell epitopes in the islets^28,29^ and peripheral blood of human T1D patients^30–32^. In the NOD mouse^33^, over 90% of the anti-insulin CD4^+^ T cell clones target amino acids 9-23 of the insulin B chain (insB:9-23)^27^. In our study, we hypothesized that T1D is caused by a gut microbiota-derived epitope via molecular mimicry mechanism and we demonstrated that the hprt4-18 peptide could activate insB:9-23 specific T-cells. Further, colonization of the female NOD mice enhanced T1D onset, increasing inflammatory cells in the spleen and pancreatic lymph nodes (PLNs). Finally, using data from the DIABIMMUNE study, we showed that children harboring the hprt4-18 sequence had a higher rate of seroconversion.

Recent studies suggest that alterations in the gut microbes and T1D could be linked to several factors, such as increased gut permeability^34^ and microbial metabolites, including SCFAs^16^ ^35^ and other proinflammatory metabolites. Here, we further investigated the impact of *P. distasonis* on the host. We focused on the gut microbiome and intestinal immune cell composition, gut permeability, cytokine levels, and serum metabolome using specific pathogen-free (SPF) NOD mice and germ-free (GF) NOD mice. Finally, to further test our molecular mimicry hypothesis, we tested whether *P. distasonis* lysate could stimulate insB:9-23 specific T-cells.

## 2. Materials and Methods

### 2.1 Animals

NOD/ShiLtJ mice were purchased from the Jackson Laboratory facility. Mice were maintained and bred in the Boston College Animal Care Facility. The mice were housed in specific pathogen-free conditions with unrestricted access to autoclaved water and bedding in a 12-h dark/light cycle. All the animal experiments were conducted as per the regulations and ethics guidelines of the National Institute of Health (NIH) and were approved by the Institutional Animal Care and Use Committee (IACUC) of Boston College (Protocol No.#B2019-003, B2022-006, 2019-004 and 2022-010). Mice were weaned at 3-weeks of age.

NOD GF mice were housed in sterile conditions utilizing Trexler-type plastic isolators. Mice were exposed to a 12:12-hour light-dark cycle and provided with autoclaved tap water and irradiated sterile pellet (breeding diet: Altromin 1414, Altromin, Germany) ad libitum. The sterility of NOD GF mice was verified biweekly by ensuring the absence of bacteria, molds, and yeast through aerobic and anaerobic cultivation of mouse feces and swabs from the isolators in meat-peptone broth, Sabouraud-dextrose and VL (Viande-Levure), followed by plating on blood, VL and Sabouraud agar plates^36^. The animal experiments were conducted as per the regulations and ethics guidelines approved by the Committee for Protection and Use of Experimental Animals of the Institute of Microbiology of the Czech Academy of Science, v.v.i. (approval ID: 117/2013).

### 2.2 Animal Treatment

3- week-old NOD mice, after weaning, were colonized as described previously. Briefly, the mice were orally gavaged for 4 weeks with either saline or live *P. distasonis* bacteria at the concentration of 1 ×10^8^ cfu/mouse/day. Bacterial colonization was determined at 10 weeks of age. Mice were sacrificed at 12 weeks of age and the pancreas, intestine, and serum were collected to perform further analysis.

For GF mice colonization, GF mice were orally gavaged once either with saline or *P. distasonis* bacteria at the concentration of 1 ×10^8^ cfu/mouse. Bacterial colonization was determined at 10 weeks of age. Mice were sacrificed at 12 weeks of age, and organs such as the pancreas, intestine, and serum were collected to perform further analysis.

For Trimethylamine N-oxide dihydrate (TMAO) treatment, TMAO powder was diluted in 1x PBS to make stock concentrations of 16 mg/ml and 32 mg/ml followed by filtration through 0.2 μM filters with syringes. Stocks were aliquoted and stored at −20°C and only defrosted when the mice were about to be injected. Littermate-matched NOD female mice were divided into three groups, 1. Saline, 2. Low TMAO (80 mg/kg body weight), and 3. High TMAO (160 mg/kg body weight). Mice were weighed weekly, then intraperitoneally injected with saline and TMAO-designated concentrations twice per week according to their assigned group for seven weeks. At 12 weeks of age, NOD mice were sacrificed to collect serum, pancreas, feces, spleen, kidney, liver, and intestines.

### 2.3 Histopathological sectioning and staining

The formalin-fixed pancreas was dehydrated using an ethanol gradient, followed by embedding in paraffin to perform histological and eosin staining. The pancreas was sectioned transversely at a thickness of 5 µm per section using a Leica RM2155 microtome. The paraffin sections were then stained with a hematoxylin and eosin (H&E) staining kit (Vector Laboratories). Images of the islets were captured using a Zeiss AxioImager Z2 upright microscope to determine the insulitis score. The islets were scored as follows: no insulitis, peri-insulitis, moderate insulitis, and severe insulitis. The insulitis scores were analyzed, and treatment groups were compared using an unpaired student t-test with Welch’s correction or one-way ANOVA with post-hoc test. The insulitis index was calculated to quantify the degree of inflammation and immune cell infiltration in the islets. No insulitis islets (0% infiltration) were scored as 0, peri-insulitis islets (<25% infiltration) were scored as 1, moderate insulitis islets (<50% infiltration) were scored as 2, and severe insulitis islets (>50% infiltration) were scored as 3. To calculate the insulitis index, the following formulate was used:

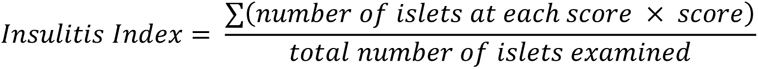

For example:

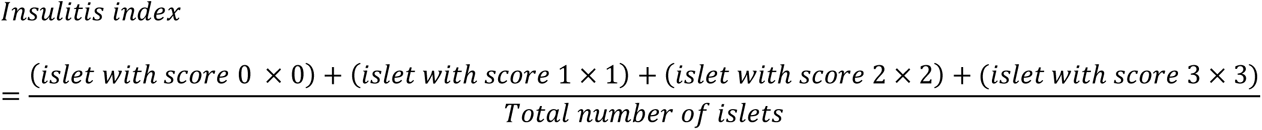

### 2.4 Intraepithelial Lymphocyte (IEL) isolation and flow cytometry

Intraepithelial lymphocyte (IEL) cells were isolated following previously established protocols^37^. Briefly, intestines were collected post-sacrifice, washed with phosphate-buffered saline (PBS), and longitudinally cut into 1-inch pieces to expose the inner epithelial layer. The intestine pieces were thoroughly washed with PBS containing 2% fetal bovine serum (FBS) to remove fecal particulates, repeating the process 2–3 times. Subsequently, the washed intestine pieces were agitated in freshly prepared 1mM dithiothreitol (DTE) solution for 20 minutes. After incubation, cells were collected from the supernatant and filtered through a 70μm filter. The DTE incubation process was repeated once more to ensure maximum cell recovery. The collected cells from both rounds of incubation were combined and further purified using a 44/67 Percoll gradient to isolate the intraepithelial lymphocyte (IEL) cells. The cell suspension was washed twice with RPMI media containing FBS before surface labeling with appropriate fluorochrome-conjugated monoclonal antibodies, as detailed in **Table S2**. The samples were analyzed using a BD FACSAria flow cytometer, and the acquired data were analyzed using FlowJo10 software. The gating strategy employed to identify the cell population is described in **Fig S2** and **Fig S3**.

### 2.5 16S rRNA gene sequencing

Fecal pellets were obtained from 10-week-old colonized NOD mice immediately snap freezed, and stored at −80°C. Stored fecal pellets were used to obtain DNA samples using a Qiagen Powerfecal pro-DNA isolation kit. 16S sequencing was performed at the TGen Integrated Microbiomics Center (TIMC). Bacterial DNA was quantitated by BactQuant assay^38^. 16S rRNA gene libraries were created by amplifying the variable region 4 (V4) using dual-index primers that included the Illumina adapters^39^. The quality of the 16S rRNA amplicon library pool was assured using Tapestation, Qubit, and KapaQuant analysis. Amplicons were pooled and sequenced on one Illumina MiSeq Nano (2 x 250 bp) run. The resulting data from the MiSeq run (MiSeq 1027) yielded 711,008 bp that successfully passed the filter, with a read range of 6,497 – 28,665 reads per sample. The average quality score (Q30) obtained was 32.9.

### 2.6 Serum metabolomic analysis

All serum samples were thawed at room temperature on a rotating plate for 20-30 minutes and then kept on ice during aliquoting. Each sample was vortexed before extracting 75μL with 450μL of cold extraction solvent (3:3:2 IPA: ACN: H_2_O at −20°C). An additional 5μL of serum from each sample was combined and extracted (1.44mL cold extraction solvent) to form a pooled QC sample. Alongside the samples, 5 QC plasma aliquots (75μL each) were extracted following the same protocol. All sample mixtures were vortexed for 5 seconds and stored overnight at −20°C. After the overnight extraction, all samples, QC plasma, and the pooled sample were centrifuged at 21,000 g for 10 minutes. The supernatant (100μL) was then transferred to glass LCMS vials for analysis. Samples were analyzed on a Sciex 5500 Triple Quadrupole Mass Spectrometer (Sciex, MA, USA) coupled to a Shimadzu NEXERA XR UPLC system (Shimadzu, Columbia, MD, USA). A two-solvent liquid gradient system consisting of 50 mM ammonium bicarbonate at pH = 9.4 (aqueous) and 100% Acetonitrile (organic) was used over 30 minutes applying Hydrophobic Interaction Liquid Chromatography (HILIC) to separate targeted metabolites. An ApHeraTM NH2 HPLC column (5μm, 15cm x 2mm) coupled to an apHeraTM NH2 HPLC guard column (5μm, 1cm x 2mm) and a pre-column filter (0.5μm A-102 frit) were used for chromatographic separation. Samples were analyzed using a scheduled MRM method with a 4-minute detection window for each metabolite transition. Resulting data was trimmed of non-detected metabolites and then statistically analyzed via MetaboAnalyst based on desired sample groupings, using: mean normalization, log transformation, and Pareto scaling.

### 2.7 Luminex assay

Luminex cytokine assay was performed utilizing the Mouse Cytokine/ Chemokine Magnetic Bead Panel (Millipore# MCYTOMAG-70K). Reagents were prepared as per kit instructions (including wash buffers, sample matrix, beads, standards, etc.). The assay was performed as per the manufacturer’s instructions. Briefly, 200 μL of Wash Buffer was added into each well and the sealed plate was shaken for 10 minutes at room temperature. Following this, the Wash Buffer was decanted, 25 μL of each Standard or Control was added to the appropriate wells, and 25 μL of Assay Buffer was added to the sample wells. For background, standards, and control wells, 25 μL of the Serum Matrix solution was added. 25 μL of diluted serum samples (1:1 in assay buffer) were added into the appropriate wells. 25 μL of premixed beads were added to each well. The plate was sealed, wrapped with foil, and incubated with agitation overnight at 2-8°C. After incubation, the plate was washed twice, and 25 μL of Detection Antibodies were added into each well and incubated with shaking for 1 hour. Next, 25 μL of Streptavidin-Phycoerythrin was added to each well containing Detection Antibodies; the plate was sealed, covered with foil, and incubated with shaking for 30 minutes. Following incubation, the plate was washed twice, and 150 μL of Sheath Fluid was added to all wells; then, the beads were resuspended on a plate shaker for 5 minutes. The plates were read using the then read on Bio-Plex®200 following manufacturers’ specifications and using Bio-Plex Manager software v6.2. The samples were analyzed for 32 analytes, including Eotaxin, G-CSF, GM-CSF, IFN-γ, IL-1α, IL-1β, IL-2, IL-3, IL-4, IL-5, IL-6, IL-7, IL-9, IL-10, IL-12 (p40), IL-12 (p70), IL-13, IL-15, IL-17, IP-10, KC, LIF, LIX, MCP-1, M-CSF, MIG, MIP-1α, MIP-1β, MIP-2, RANTES, TNF-α, and VEGF.

### 2.8 NOD T-cell hybridomas stimulation assay

The NOD T-cell hybridomas activation experiment was performed as described previously^26^. To obtain *P. distasonis* lysate, bacteria were diluted at concentrations of 10^6^, 10^7^, and 10^8^ CFU/ml in DMEM and sonicated for 10 min in a bath sonicator at room temperature. The C3g7 cell line, which expresses an abundance of MHC-II I-Ag7 (corresponding human DQ8), was used as an antigen-presenting cell (APCs). 10μM of hprt4-18 peptide (dissolved in DMSO), live *P. distasonis* at concentrations of 10^6^ and 10^7^ CFU, and *P. distasonis* lysate at concentrations of 10^5^, 10^6^, and 10^7^ were treated on 5 × 10^4^ C3g7 cells for 4 hours. Next, C3g7 cells were washed with PBS three times. 5 × 10^4^ T-cell hybridomas were co-incubated with treated C3g7 cells for another 20 hours. At the end of the experiment, the supernatant and cells were collected from each well. The supernatant was used to determine the IL-2 secretion using the IL2 Elisa kit (#BioLegend), and the cells were used to determine the protein concentration using BCA assay (#Thermo) for normalization.

### 2.9 Bioinformatic analysis and statistics

Bacterial 16S rRNA amplicon sequencing reads were filtered and trimmed to assess microbial diversity and perform statistical analysis. Dada2^40^ was used to convert amplicon sequences into an Amplicon Sequence Variant (ASV) table using the Ribosomal Database Project Training Set^41^.

The R (version 4.1.2) packages Phyloseq^42^ and vegan were employed for exploratory and inferential analyses, including alpha and beta diversity estimates, non-metric multidimensional scaling (NMDS) analysis using Bray–Curtis dissimilarity, Principal Components Analysis (PCA), and taxa agglomeration. Statistical significance for alpha diversity was assessed using ANOVA, while PERMANOVA was used for Bray-Curtis dissimilarity. Differential ASV abundance was evaluated at each time point using edgeR^43^ with two-sided empirical Bayes quasi-likelihood F-tests. P-values were corrected using the Benjamini-Hochberg false discovery rate (FDR), with FDR < 0.05 considered statistically significant^44^.

### 2.10 Statistics

Statistical analysis was conducted using the unpaired Student’s t-test to compare two groups. Significance levels were denoted as *p < 0.05, **p < 0.01, and ***p < 0.001.

For insulitis scores, either an unpaired Student’s t-test or a one-way analysis of variance (ANOVA) with Tukey’s post-hoc test was performed to compare multiple groups. Significance was determined at p < 0.05. All statistical analyses were conducted using GraphPad Prism Version 9.0 unless otherwise specified in the figure legends.

## 3.0 Results

### 3.1 *P. distasonis* colonization has a limited impact on gut microbiome composition in female NOD mice

To examine the impact of *P. distasonis*, we first focused on the gut microbiome composition after the colonization. To this end, we used fecal samples from 10-week-old female NOD mice, which were either orally gavaged with *P. distasonis* or saline for four weeks (starting with 3-week-old mice). *P. distasonis* colonization did not affect alpha diversity, indicating that overall species richness remains stable **(Fig1A and Fig S1A-C)**. On the other hand, the beta diversity was significantly altered by the colonization, indicating an alteration in the microbiome composition (p=0.016) **(Fig 1B)**. We also observed no significant differences in the phylum **(Fig 1C)**, family **(Fig 1D)**, class and order levels **(Fig S1D & S1E)**, as well as the genus **(Fig 1E)** level. In total, we identified 188 amplicon sequencing variants (ASVs) **(Table S1)**, with 17 ASVs showing increased abundance and 11 ASVs showing decreased abundance following *P. distasonis* colonization (p <0.05). Interestingly, most significant alterations within the annotated 28 ASVs occurred in members of *Lachnospiraceae* family. However, the alteration was not unidirectional; while ASV6, ASV7, and ASV37 were decreased, ASV36, ASV85, and ASV66 were increased **(Table S1 and Fig 1F)**. Overall, these results indicate that *P. distasonis* colonization had a limited impact on the gut microbiome composition.

**Fig 1:**
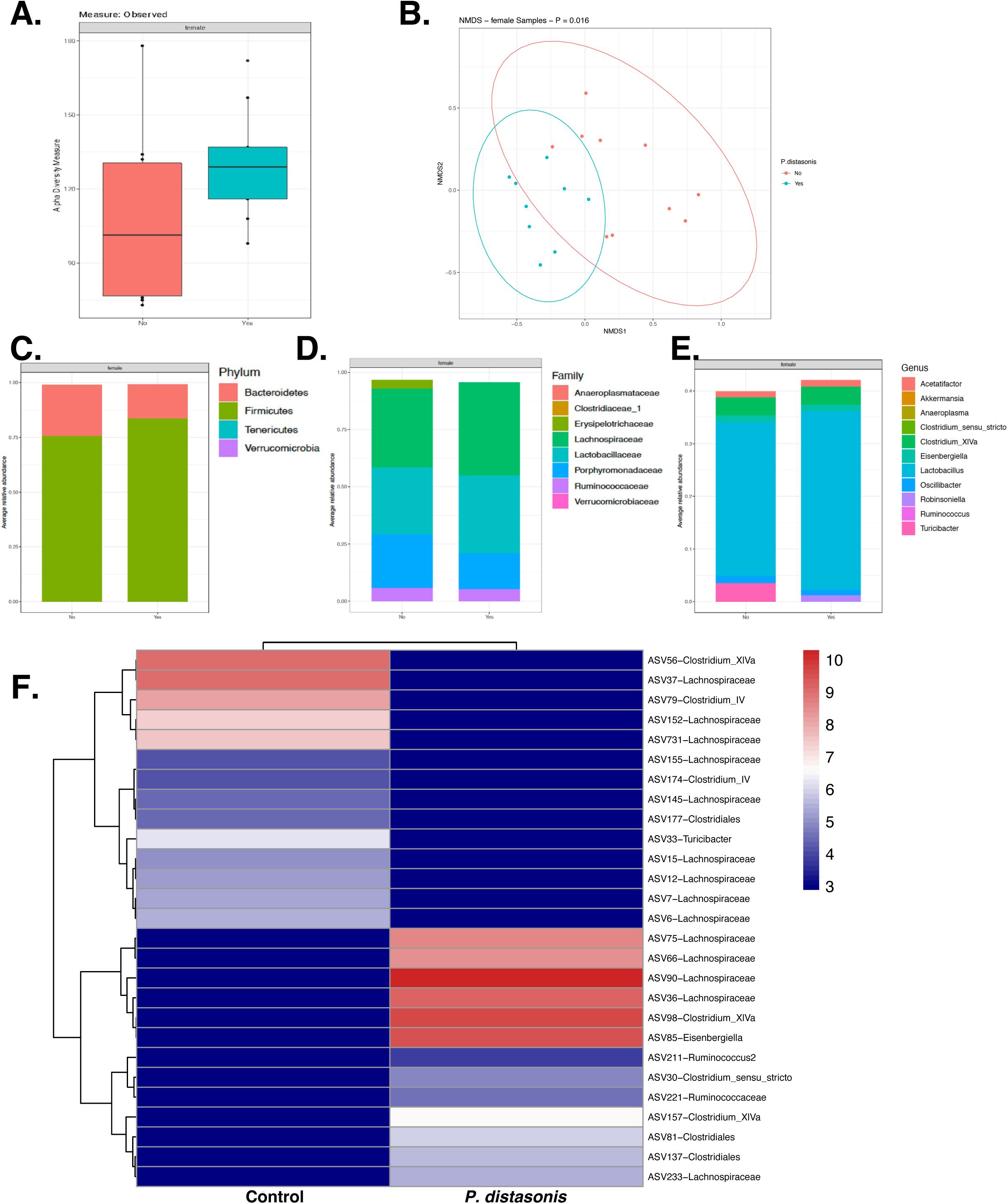
The effect of *P. distasonis* on gut microbiome profile of NOD female mice. **(A)** Alpha diversity index **(B)** Beta diversity **(C)** Average relative abundance of bacterial phylum **(D)** family and **(E)** genera. Statistical analysis was performed with the Benjamini and Hochberg method using two-tailed t-tests to control the false discovery rate (FDR). **(F)** Heat map showing the relative abundance of the ASVs significantly different between *P. distasonis* colonized and saline-gavaged mice. Each column represents the mean of each group, and each row represents an ASV.

### 3.2 *P. distasonis* decreases inflammatory immune cells in small intestines without altering gut permeability

We previously reported that *P. distasonis* colonized NOD female mice, had an increase in CD8+ T-cells, F4/80+ macrophages, and dendritic cells, along with a decrease in FoxP3+ regulatory T cells in splenocytes. Similar alterations were also observed in the pancreatic lymph nodes for FoxP3+ T-regulatory cells and macrophages. However, the effect of *P. distasonis* on systemic inflammation and intestinal immune cell composition remained elusive. To determine whether the elevated incidence of T1D is associated with increased inflammation in the gut, we examined the direct effects of *P. distasonis* in the intestines, where the bacterium directly interacts with the immune system. We initially assessed the intestinal intraepithelial lymphocytes (IELs) composition in 12-weeks old female NOD mice.

Flow cytometry data revealed a 2.4-fold decrease in CD4+ T-cells **(Fig. 2A)** and a 2.6-fold decrease in B-cells **(Fig. 2B)** in *P. distasonis* colonized mice. However, there was no significant difference in total T-cells, CD4+CD25+ T-cells, and CD8+ T-cells **(Fig. S2C-E)**. Additionally, we examined the innate cell composition in IELs, including dendritic cells, macrophages (total, resident, and circulatory macrophages), and eosinophils. We observed a 1.7-fold decrease in resident macrophages **(Fig. 2C)** in *P. distasonis* colonized mice, while no significant differences were observed in dendritic cells, total macrophages, circulatory macrophages, and eosinophils populations **(Fig. S3B-F)**. Further analysis of CD4+ and CD8+ T-cell subsets showed a 1.76-fold decrease in CD4+ T-effector and a 2.1-fold decrease in CD4+ T-central cell populations **(Fig. 2D)**. We also observed a 3-fold decrease in CD8+ T-central memory cell population upon *P. distasonis* colonization in IELs **(Fig. 2E)**. Overall, *P. distasonis* colonization reduced CD4+ effector T-cells, both CD4+ and CD8+ central memory T-cells, B-cells, and resident macrophage populations. These results suggest that *P. distasonis* does not stimulate any inflammatory immune cell population in the small intestine upon colonization; in contrast, colonization creates a less inflammatory intestinal environment.

**Fig 2:**
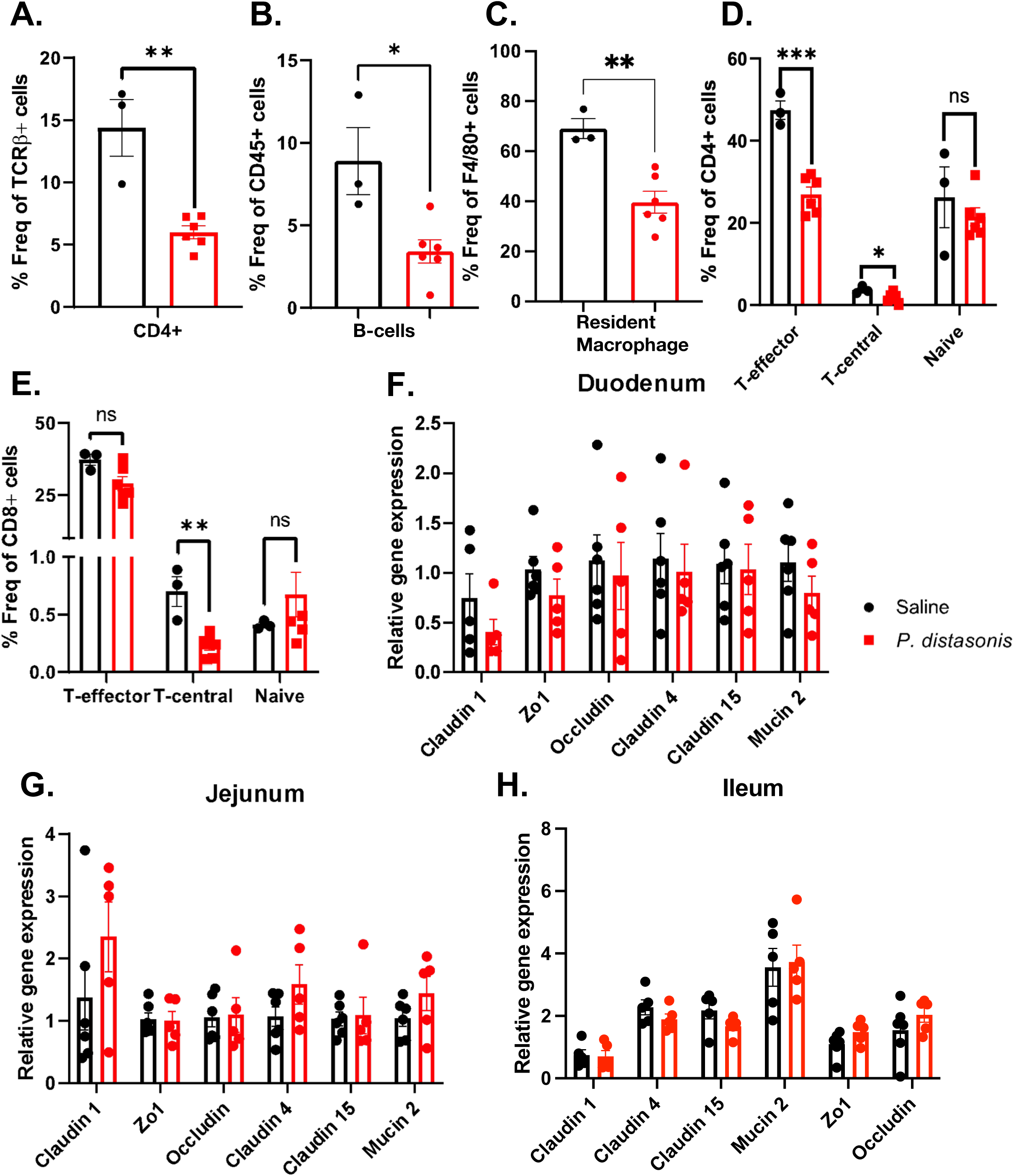
The effect of *P. distasonis* colonization on intestinal inflammation in NOD female mice. **(A)** CD4+ cells as a percentage of TCR-β+ CD45+ immune cell subsets. **(B)** Percentage of B-cells in CD45+ cells subsets. **(C)** Percentage of CD11b+ (dendritic cells subsets in CD45+ immune cells. **(D)** CD44^hi^CD62L^lo^ (T_EM_), CD44^hi^ CD62L^hi^ (T_CM_), CD44^lo^ CD62L^hi^ (Naive) T-cells in CD4+ T-cells. **(E)** CD44^hi^CD62L^lo^ (T_EM_), CD44^hi^ CD62L^hi^ (T_CM_), CD44^lo^ CD62L^hi^ (Naive) in CD8+ T-cells subsets population in saline and *P. distasonis*–gavaged mice. Relative gene expression of gut permeability-related genes in **(F)** duodenum, **(G)** jejunum, **(H)** ileum in *P. distasonis* colonized NOD mice compared to saline NOD mice (n=5/group). Data were expressed as mean ± SEM. *p<0.05, **p <0.01, ***p <0.0001. Statistical analysis was performed using the two-tailed unpaired Student’s t-test.

To determine the impact of *P. distasonis* colonization on the systemic immune response, we employed a Luminex assay analyzing 32 different cytokines, chemokines, and growth factors on serum samples from 12-week-old *P. distasonis*-colonized female NOD mice compared to control mice (n=8-9/group). From those 32 analytes, only IL-15 levels were 1.9-fold decreased upon *P. distasonis* colonization **(Fig. S4)**. Altogether, these results are consistent with our observations in the intestines and indicate that *P. distasonis* does not stimulate a non-specific, pro-inflammatory, or anti-inflammatory cytokine response that can explain increased insulitis or diabetes rates of NOD female mice upon colonization. Subsequently, we investigated the impact of *P. distasonis* on gut permeability as another potential mechanism that might stimulate inflammation and contribute to diabetes acceleration. Gene expression analysis of claudin family proteins (claudin1, claudin4, and claudin15) and tight junction proteins (ZO1, Occludin, Mucin2) in the duodenum, jejunum, and ileum did not reveal significant changes upon *P. distasonis* colonization **(Fig. 2F-H)**. Taken together, *P. distasonis* colonization doesn’t increase gut permeability in the small intestines.

### 3.3 *P. distasonis* does not alter serum metabolome composition

While there were only 28 ASVs altered by the *P. distasonis* colonization, we observed a significant alteration in beta diversity. To determine whether these alterations have an impact on the serum metabolome, we employed, targeted metabolomics using serum samples from 12-week-old female NOD mice. In total, 255 metabolites were determined in this analysis. The principal component analysis (PCA) did not reveal any significant differences between the groups **(Fig. 3A)**. Among all 255 targeted metabolites, we identified no significant change in any of the metabolites between *P. distasonis* and the saline-treated group **(Table S3, Fig 3B-C)**. However, TMAO, a metabolite directly related to the gut microbiome and inflammation tended to increase but it was not a significant increase **(Table S3**, **Fig. 3B)**. Overall, these findings suggest that *P. distasonis* colonization has no impact on the serum metabolome composition.

**Fig 3:**
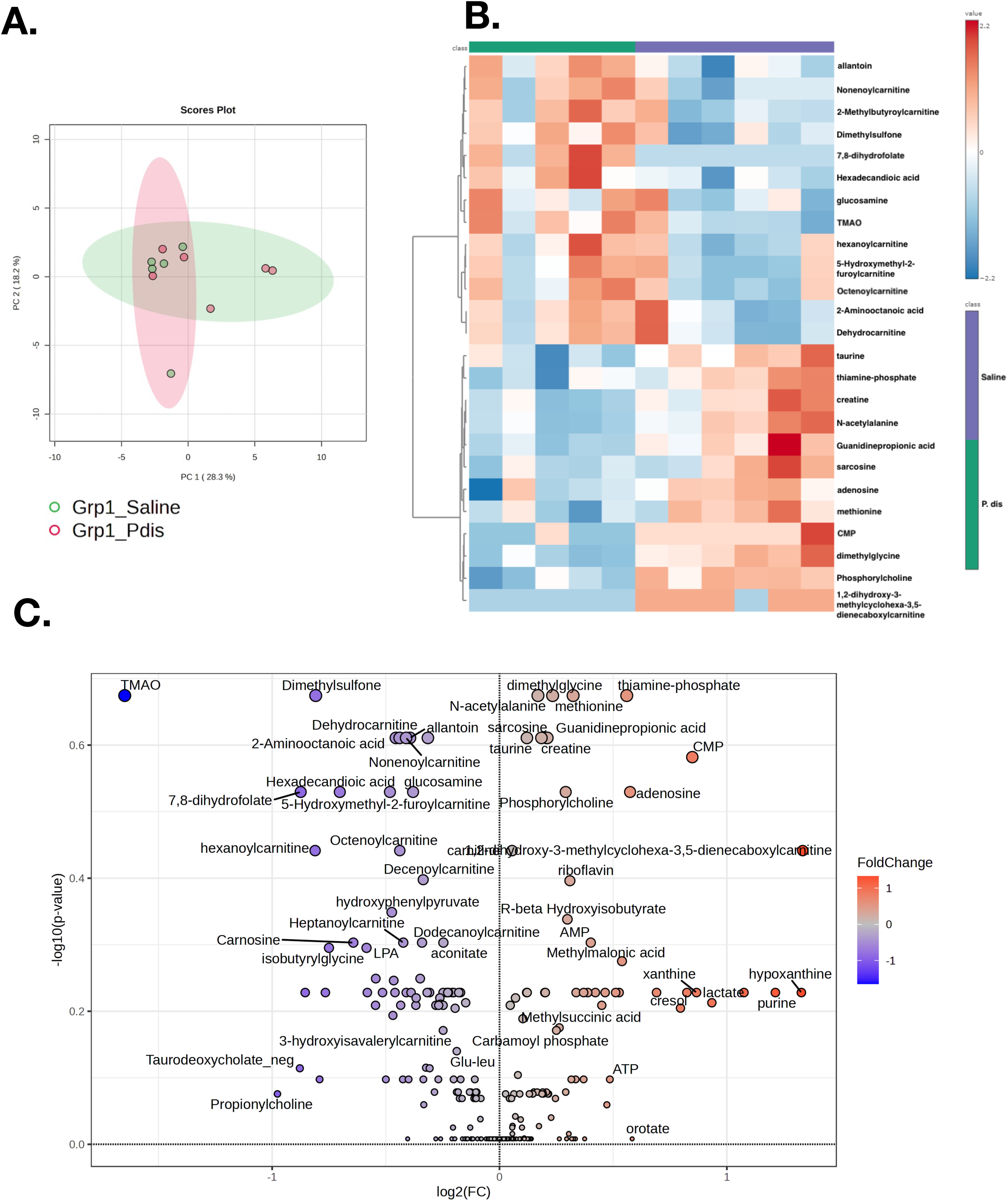
The effect of *P. distasonis* colonization on serum metabolites composition in NOD female mice. (A) Principal Coordinate Analysis (PCA) plot showing serum metabolites comparison in saline and *P. distasonis* colonized SPF female NOD mice. **(B)** Heatmap showing 25 of the most altered metabolites between saline and *P. distasonis* oral gavaged mice. Each column represents an individual mouse, and each row represents a metabolite. **(C)** Volcano plot of serum metabolites with fold change threshold (|log2 (FC)|>1.2) and t-tests threshold (-log10(p)>0.1). The red dots represent metabolites above the threshold. Fold changes are log2 transformed, and p-values are log10 transformed of saline (n=6) and *P. distasonis* (n=5) oral gavaged NOD female mice.

### 3.4 TMAO does not alter insulitis in NOD female mice

TMAO is one of the most studied inflammatory microbial metabolites and has been associated with cardiovascular and metabolic diseases^45–49^. To determine whether increased diabetes rates in *P. distasonis* colonized SPF NOD mice are linked to the trend of increase in the TMAO, we decided to examine its effect on diabetes onset. To this end, we intraperitoneally injected female NOD mice (n=5-6 mice/group) either with saline (control group), 80mg/kg TMAO (low dose), or 160mg/kg TMAO (high TMAO group) **(Fig. 4A)**. Mice were weighed weekly and there were no significant differences in body weight upon TMAO administration **(Fig. 4B)**. After seven weeks of consecutive injections, 12-week-old, prediabetic mice were sacrificed to determine the insulitis scores. We demonstrated that there was no significant difference in the insulitis scores between the control mice and the TMAO groups (85-113 islets per group, **Fig. 4C**). We further calculated the insulitis index scores **(Fig. 4D)**, however, once again, we did not identify any significant differences.

**Fig 4:**
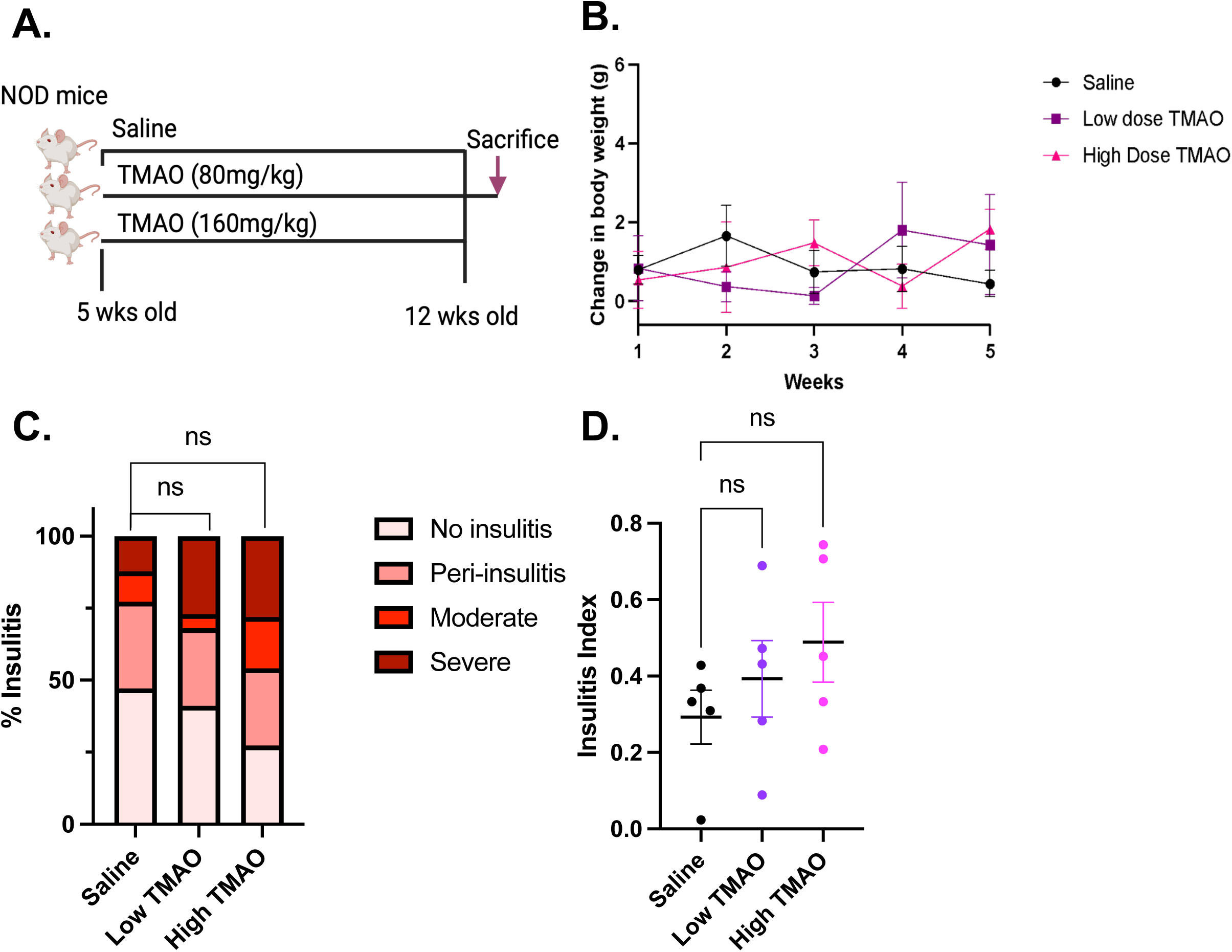
The effect of TMAO on NOD female mice insulitis. **(A)** Schematic overview of the low (80mg/kg) and high Dose (160mg/kg) TMAO-injected NOD female mice experiment (n =5-6/group). 5-week-old NOD mice were injected with different doses of TMAO for seven weeks. **(B)** Change in body weight per week. **(C)** Percentage of insulitis score in low and high-dose treated mice compared to saline-treated mice. Statistical analysis was performed using the one-way ANOVA.

### 3.5 *P. distasonis* accelerates insulitis in NOD germ-free (GF) female mice

To determine the isolated impact of *P. distasonis* colonization on insulitis and mitigate potential influences from other gut microbes in specific pathogen-free (SPF) NOD mice, we utilized female GF NOD mice. This approach aimed to eliminate any confounding factors stemming from the presence of other gut microbes. The GF female NOD mice (n=5-6) were orally gavaged once with *P. distasonis* after weaning **(Fig. 5A)**. Colonization was confirmed in 10-week-old GF NOD mice using qPCR **(Fig. 5B)**. The mice were sacrificed at 12 weeks of age to determine the insulitis scores. Consistent with our previous study in SPF NOD mice, *P. distasonis* colonization alone increased insulitis in GF NOD mice **(Fig. 5C)**. It caused a 2.8-fold increase in severe insulitis in islets and a 5.6-fold decrease in islets with no insulitis compared to the control animals. This data indicates that *P. distasonis*, independent of other gut microbes, can induce insulitis in NOD mice. To investigate the mechanism of *P. distasonis*-induced insulitis and its potential link to gut permeability, we assessed the expression of gut permeability associated genes. We assessed the gene expression of claudin 1, claudin 4, ZO1, claudin 15, mucin2, and occludin in different parts of the small intestine. Similar to our findings in SPF mice, we did not observe any differences in the duodenum **(Fig. 5D)**, jejunum **(Fig. 5E)**, and ileum **(Fig. 5F)** indicating that *P. distasonis* colonization does not alter gut permeability.

**Fig 5:**
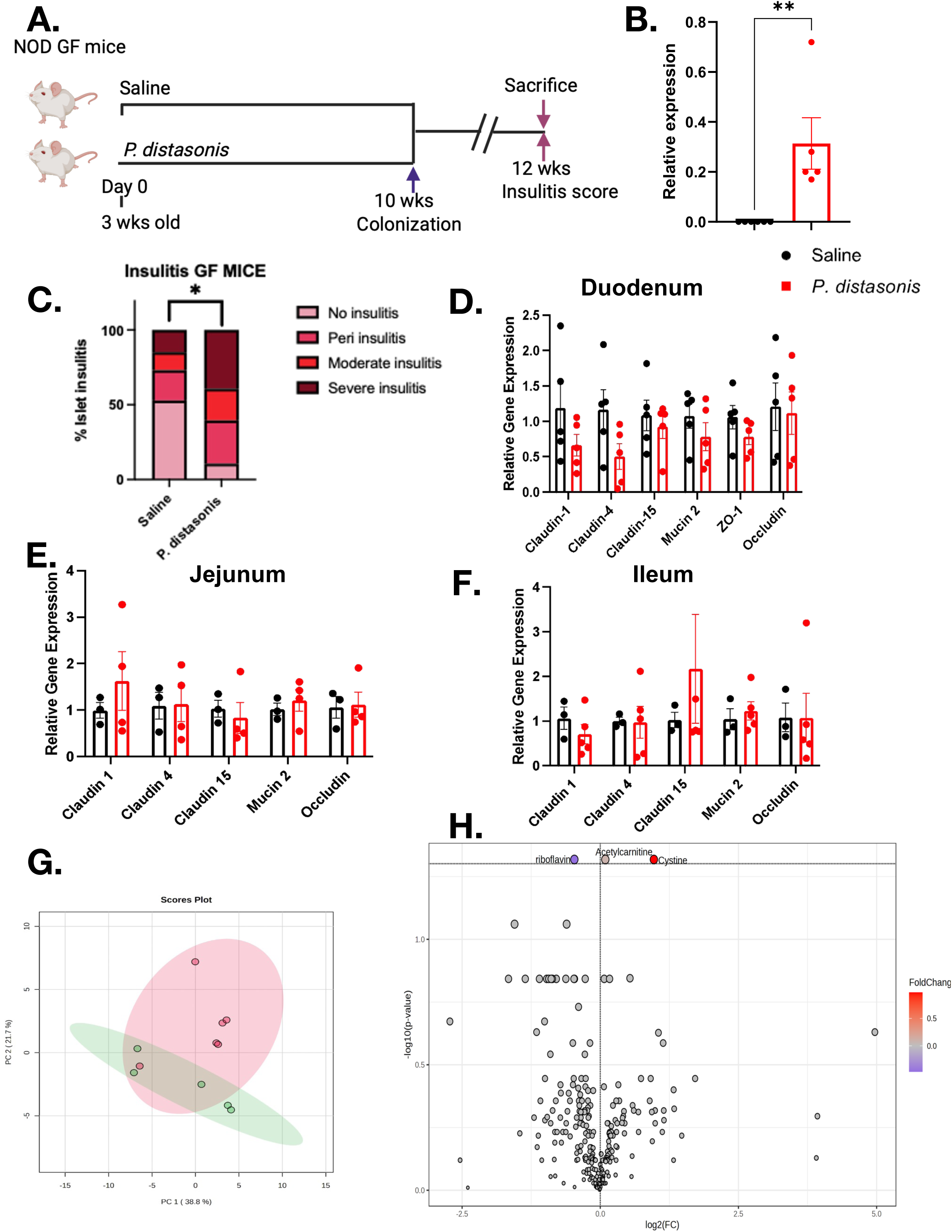
The effect of *P. distasonis* colonization on insulitis, gut permeability, and serum metabolome in germ-free (GF) NOD female mice. **(A)** Schematic overview of the *P. distasonis* oral gavage experiments (n = 13, saline, n=11, *P. distasonis*). The time point of week 10 represents the fecal sample collection for qPCR colonization experiment for *P. distasonis* colonization, and the red arrow shows the time point (week 12) for pancreata collection for insulitis analysis (n = 5 mice/group). **(B)** Quantification of insulitis scores from *P. distasonis* colonized, or saline-gavaged female NOD GF mice at week 12 (n = 5/group, n = 150 to 180 islets/group). **(C)** Relative gene expression of gut permeability-related genes in duodenum, **(D)** jejunum, **(E)** ileum in *P. distasonis* colonized NOD GF Female mice compared to saline NOD GF mice (n=3-5/group). Data were expressed as mean ± SEM. *p<0.05, **p <0.01,***p <0.0001. Statistical analysis was performed using an unpaired Student’s t-test for insulitis and gene expression analysis. **(F)** Principal Coordinate Analysis (PCA) plot showing a comparison of serum metabolites between saline and *P. distasonis* oral-gavaged GF female NOD mice. **(G)** Heatmap showing 25 of the most altered metabolites between saline and *P. distasonis* oral-gavaged GF female NOD mice. Each column represents an individual mouse, and each row represents a metabolite. **(C)** Volcano plot of serum metabolites with fold change threshold (|log2 (FC)|>1.2) and t-tests threshold (-log10(p)>0.1). The red dots represent metabolites above the threshold. Fold changes are log2transformed, and p values are log10 transformed of saline (n=5) and *P. distasonis* (n=6) oral gavaged NOD GF female mice.

### 3.6 *P. distasonis* colonization does not alter serum metabolite composition in GF NOD mice

To determine the direct impact of *P. distasonis* on metabolome, we performed targeted metabolite analysis using serum samples obtained from the GF mice. In total, we identified 255 metabolites **(Table S4)** and among them, only three metabolites were significantly altered. However, the magnitude of these changes was relatively low. The PCA did not identify significant differences between groups **(Fig. 5G)**. Specifically, cystine levels increased by 1.9-fold, and acetylcarnitine levels increased only by 1.06-fold, while riboflavin levels decreased by 1.4-fold in *P. distasonis* colonized mice **(Fig. 5H)**. Overall, our findings are consistent with our SPF findings and indicate that *P. distasonis* has minimal impact on serum metabolome composition in GF mice.

### 3.7 *P. distasonis* lysate activates insB:9-23 specific T-cells hybridomas

In this study, we examined different effects of *P. distasonis* colonization on SPF NOD mice and GF NOD mice to identify potential other factors that could affect diabetes onset stimulated by *P. distasonis*. However, we could not identify any factors including gut permeability, a non-specific inflammatory immune response in the gut, or significant alterations in the metabolome, or any effect related to TMAO. These observations reaffirm our original hypothesis that *P. distasonis* stimulates diabetes onset via molecular mimicry. In our previous study, we used chemically synthesized 15-amino acid long, hprt4-18, and insB:9-23 peptides and showed that hprt4-18 can stimulate both insB:9-23 specific human T-cell clones and NOD mice T-cell hybridomas^26^. Nonetheless, it was previously unknown whether APCs could effectively process the whole bacterial lysate and present hprt4-18. To test this, we treated APCs either with live *P. distasonis* or *P. distasonis* lysate. After 4 hours of treatment, APCs were co-cultured with insB:9-23 specific IIT-3 T-cell hybridomas, and T-cell activation was assessed by measuring IL-2 secretion. hprt4-18 peptide (10 µM) used as a positive control, successfully activated the T-cell hybridomas (**Fig. 6**). Notably, APCs were also able to process *P. distasonis* lysate (CFUs, 10^5^ and 10^6^) and activate these insB:9-23 specific T-cells. The live *P. distasonis* did not stimulate the T-cells potentially because hprt is a cytoplasmic protein. These findings support our molecular mimicry hypothesis and show that the APCs can effectively process *P. distasonis* proteins, specifically the hprt protein, and stimulate insB:9-23 specific T-cell hybridomas (**Fig 6**).

**Fig 6:**
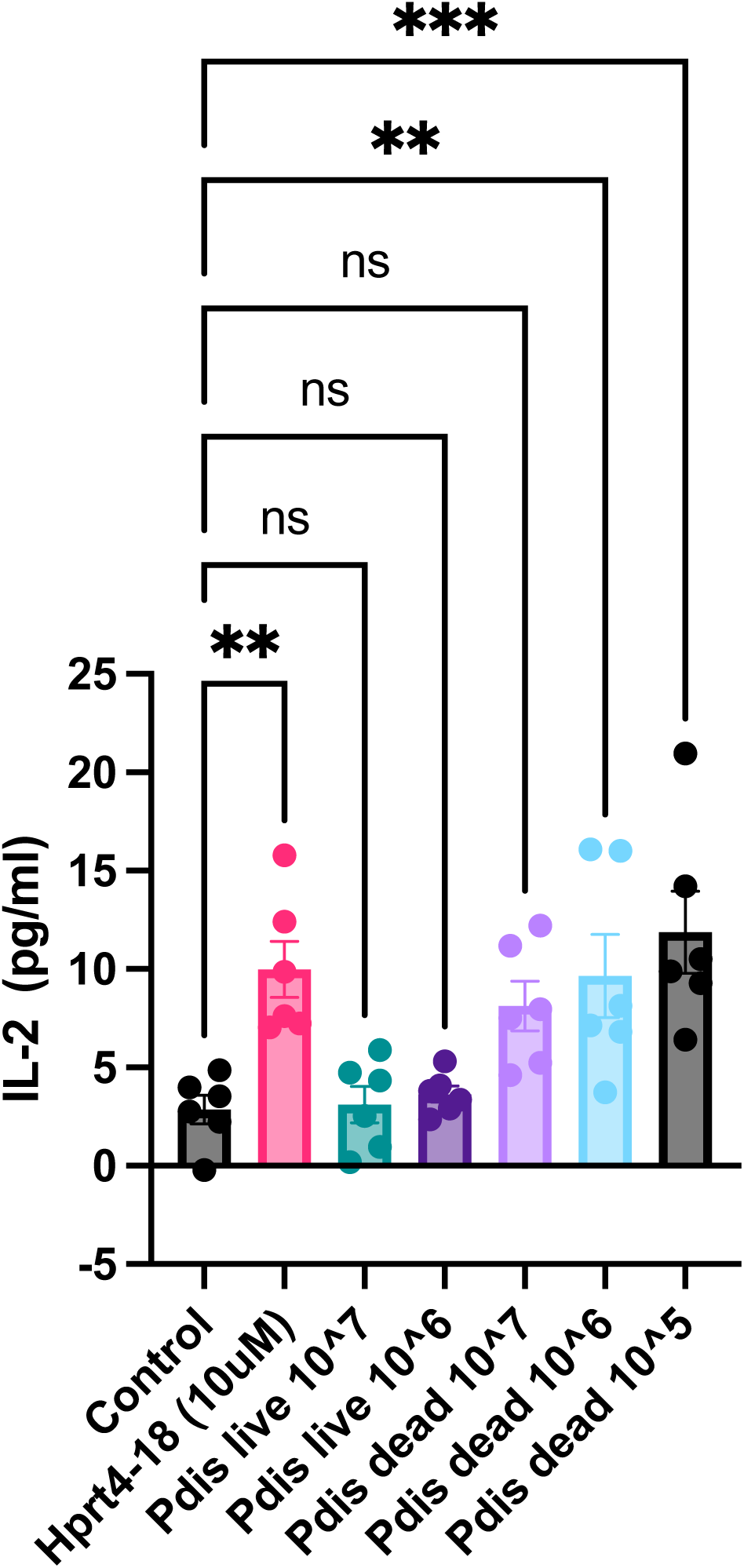
The effect of *P. distasonis* on insB:9-23 specific NOD Mice T-cells hybridomas. IIT-3 hybridomas response to DMSO (control), hprt4-18 peptide, *P. distasonis* live or dead (10^7^, 10^6^, and 10^5^ CFU) where peptides/live or dead bacteria were presented to hybridomas as covalently linked to I-A^g7^ expressed on macrophages (C3g7 cells). Data are presented as IL-2 concentration measured using ELISA. Data were expressed as mean ± SEM. *p<0.05, **p <0.01,***p <0.0001. Statistical analysis was performed using one-way ANOVA.

## 4.0 Discussion

T1D is one of the oldest chronic autoimmune diseases, and it is still not curable. Although we have better tools to manage the disease, it still decreases life expectancy by an average of 9.9 years^50^. We cannot prevent new cases because the trigger of T1D autoimmunity is unknown. Therefore, there is an urgent need to identify environmental factors contributing to T1D onset. In this study, we examined the involvement of *P. distasonis* in T1D pathogenesis. As previously mentioned, the TEDDY and INNODIA studies reported that *Parabacteroides* is the most significantly associated genus with T1D onset^15^.

In our previous study, we also showed that the *P. distasonis* D13 strain colonization accelerates T1D onset in female NOD mice. To further define the impact of *P. distasonis* colonization on the host, we completed this comprehensive study. Our results on gut microbiome composition indicated a significant change in beta diversity, with 28 differentially abundant ASVs, mostly members of the *Lachnospiraceae* family. Interestingly, *Clostridium* species are found in this family and some of them including *Clostridium asparagiforme, Clostridium hathewayi, Clostridium sporogenes* are TMA producers^45^. This might explain the trend of increasing TMAO levels in the serum metabolome.

When we evaluated the effects of colonization on immune cell composition in the IELs, we observed a decrease in residential macrophages, B-cells, and CD4+ cells. Notably, there was also a reduction in CD4+ T-effector, T-central subsets, and CD8+ T-central subsets. Intestinal CD4+ T cells are a major population in IELs, crucial for maintaining host protective and homeostatic responses to gut microbes. However, an accumulation of CD4+ T cells, particularly the CD4+ T-effector subtype, is a hallmark of inflammation and inflammatory bowel disease (IBD)^51^ ^52^ ^53^. Similarly, B-cells play a significant role in increasing gut inflammation and villous atrophy in celiac disease^54^. Reduction in these immune cells on colonization indicates a non-inflammatory phenotype.

*P. distasonis* significantly reduced intestinal inflammation in murine models of acute and chronic colitis^55^ induced by dextran sulphate sodium (DSS) in BALB/c mice. This anti-inflammatory effect is mediated by a decrease in proinflammatory cytokines and stabilization of the intestinal microbiota ^56^. In our study, we also observed a decrease in serum IL-15 concentration, consistent with our observations in the IELs. One of the potential mechanisms explaining this anti-inflammatory function is related to having a unique surface layer that breaks down complex polysaccharides and helps them blend with intestinal tissue^57^. These findings on reduced intestinal inflammation align well with our results. Interestingly, treating high-fat diet-fed and ob/ob mice, two different models of Type 2 Diabetes and obesity, with *P. distasonis* CGMCC1.30169 reduced weight gain, hyperglycemia, and hepatic steatosis by activating intestinal gluconeogenesis and FXR pathways^58^. The authors explained this phenotype with succinate production by the bacterium. However, we did not identify an increase in succinate either in SPF NOD or in GF NOD mouse models.

There are also controversial findings indicating potential pathogenic effects of the bacterium. For example, the stools of Crohn’s disease patients repeatedly contained *P. distasonis* bacteria^59^. Likewise, in a DSS-induced colitis model, there was a correlation between *P. distasonis* abundance and severity of colitis^60^. *P. distasonis* inoculation increased inflammation in mice that already have Crohn’s disease^61^. In a similar study, *P. distasonis* weakened the gut barrier and triggered inflammation, suggesting a link to IBD^62^. In addition, *P. distasonis* produces an enzyme that can inactivate antimicrobial peptides, including β-defensin 2, keratin-derived antimicrobial peptides (KAMPs), and human neutrophil peptide 3^63^. The differences observed in all these studies may be attributed to the variety of disease models, strains, and animal models tested.

Overall, our data indicate that *P. distasonis* colonization does not create a pro-inflammatory environment in the intestine of NOD mice. In contrast, it causes a reduction in the immune cells indicating a potential anti-inflammatory effect in the gut. The underlying mechanism behind the decrease or migration of the immune cells from the intestine still needs to be determined.

The antigen presentation assay revealed that *P. distasonis* lysate can activate insB:9-23 specific T-cell hybridomas. This indicates that the APCs processing *P. distasonis* lysate proteins are capable of activating T-cells, potentially leading to anti-insulin autoimmunity. The molecular mimicry mechanism is based on the degeneracy of T-cell recognition^64^ ^65^ and can be either pathogenic^66^ or protective^67^. While molecular mimicry has long been postulated as a potential factor in autoimmune diseases^68–70^, including T1D^71–74^ (coxsackie B and rubella viruses), progress was hindered due to a lack of genomic sequences of the potential microbial proteins that might trigger this response. In our previous study, we took advantage of the growing genome databases for microbes, including growing microbiome datasets, and identified hprt4-18^75^. We showed its potential rolling T1D pathophysiology establishing cross-reactivity and determining the enhanced T1D in *P. distasonis* colonized NOD mice. Our findings reported here could not identify any other potential mechanisms that might stimulate T1D. Therefore, we decided to test our original molecular mimicry mechanism with a key missing experiment, antigen presentation. Indeed, we showed that APCs can process *P. distasonis* proteins and present and activate insB:9-23 T-cells. However, the specific role of hprt4-18 still needs to be determined in future studies using mutation models. Lumen presents a high challenge for microbial survival. Despite the evolutionary adaptations of gut microbes to colonize the gut, cell death and cell lysis remain inevitable. These findings support our original hypothesis and indicate that lysed *P. distasonis* may be processed by the APCs in the GI tract. In addition to our findings, molecular mimicry was recently linked to other autoimmune diseases including multiple sclerosis (MS). Recent studies identified pathogenic and cross-reactive antibodies as a potential link between Epstein–Barr virus (EBV) and MS onset. Researchers showed the cross-reactivity between EBV nuclear antigen EBNA1 and glial cell adhesion protein in the central nervous system^76^. Epidemiological evidence supports this link between EBV infection and MS autoimmunity^77^.

While further studies are needed to establish a causal link between human T1D and *P. distasonis*, we believe that our findings serve as a proof-of-concept study, shedding light on a potential link between gut microbiota-derived neo-epitopes and autoimmunity. While our study focused on *P. distasonis*, other pathobionts in the gut may also generate insB:9-23 specific T-cells. Furthermore, hundreds of epitopes have been identified in the pathogenesis of T1D^78^ ^79^, with the potential for T-cells to be stimulated by several gut microbiota-derived mimic epitopes. Identifying these commensal microbes could provide a better understanding of gut-immune interactions and reveal mechanisms underlying autoimmune diseases such as T1D. Identification of causal factors will guide us in developing novel therapeutic tools to prevent, cure, and manage the disease.

## Acknowledgments

We thank Prof. Emil Unanue, Xiaoxiao Wan, and Dr. Anthony Vomund (University of Washington, St. Louis) for providing insB:9-23 specific T-cell hybridomas and C3g7 APCs cells. Thanks to Patrick Autissier and Bret Judson for their assistance with flow cytometry and imaging at the Microscopy Core at Boston College, respectively. Special thanks to Dr. Babak Momeni (Boston College) for sharing his laboratory’s anaerobic chamber. We are grateful to Nancy McGilloway and Todd Gaines for their support in the Boston College Animal Care Facility. We thank our students, Lukas Rhodes and Maria Jove, for their assistance with animal maintenance. This work was supported by The G. Harold and Leila Y. Mathers Charitable Foundation Research Grant No. MF-1905-00311, a JDRF Foundation grant (K 98-99D-12813-01A), the Beatson Foundation (2023-003), and a Diabetes Research Connect grant to KG.

## Contributions

K.G. and E.A. designed the research, analyzed the data, and wrote the paper. E.A oversaw the project. Y.D.D. assisted with the bioinformatic analysis of the 16S data. K.G., C.H., and A.P. assisted with all animal experiments. K.G. performed the FACS staining and analysis, while A.R. assisted with TMAO experiment and M.S., P.S, U.K.G, and T.H. assisted with GF mice experiments and maintenance. M.K. and J.H. conducted the serum metabolomic analysis.

## Consent for publication

Not applicable.

---

**Fig S1:**
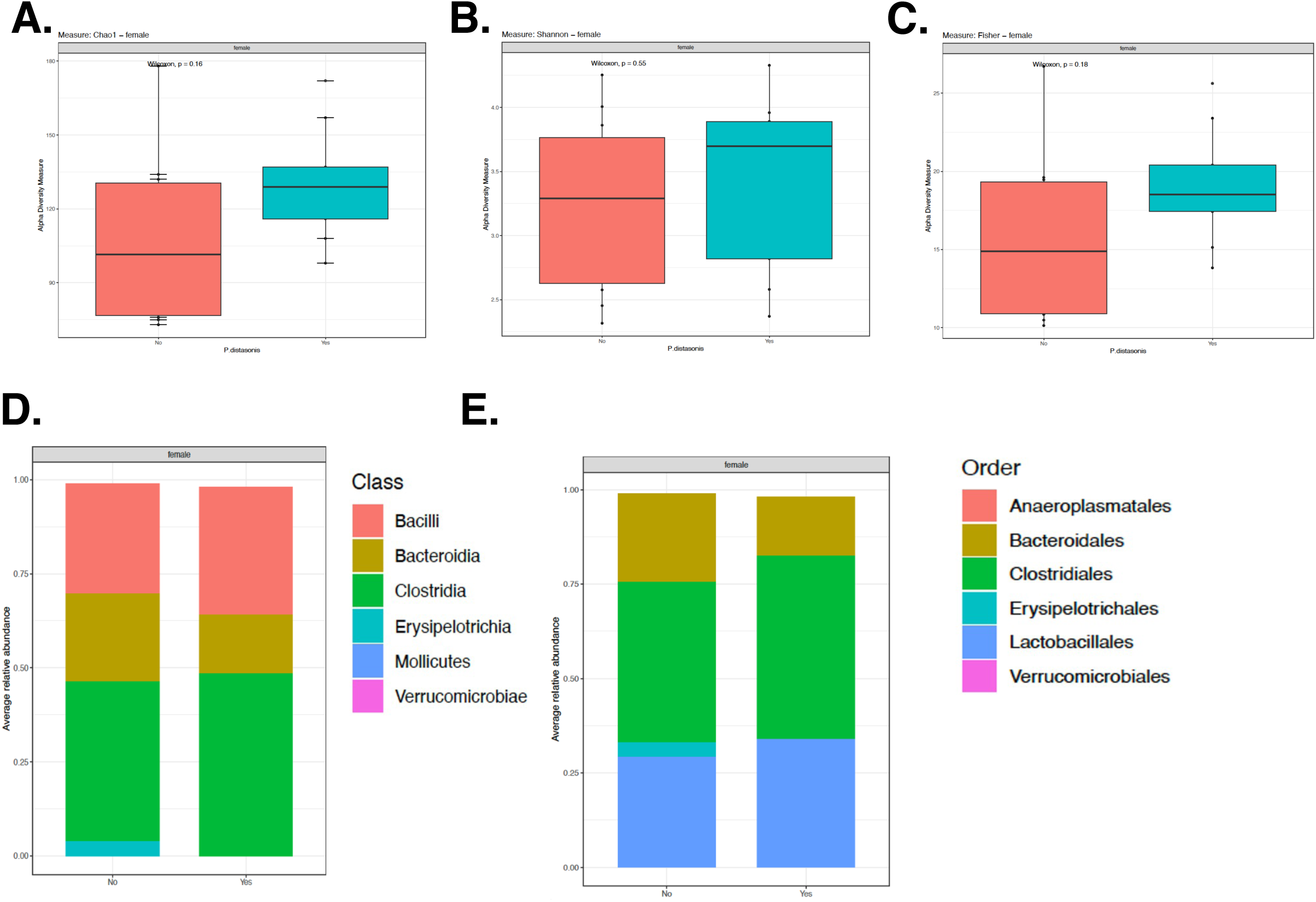
Alpha diversity indexes and Relative abundance of gut bacterium upon *P. distasonis* colonization. **(A)** Alpha diversity Chao index **(B)** Shannon index **(C)** Fisher index **(D)** Average relative abundance of bacterial class **(E)** Order between *P. distasonis* colonized and saline gavaged mice.

**Fig S2:**
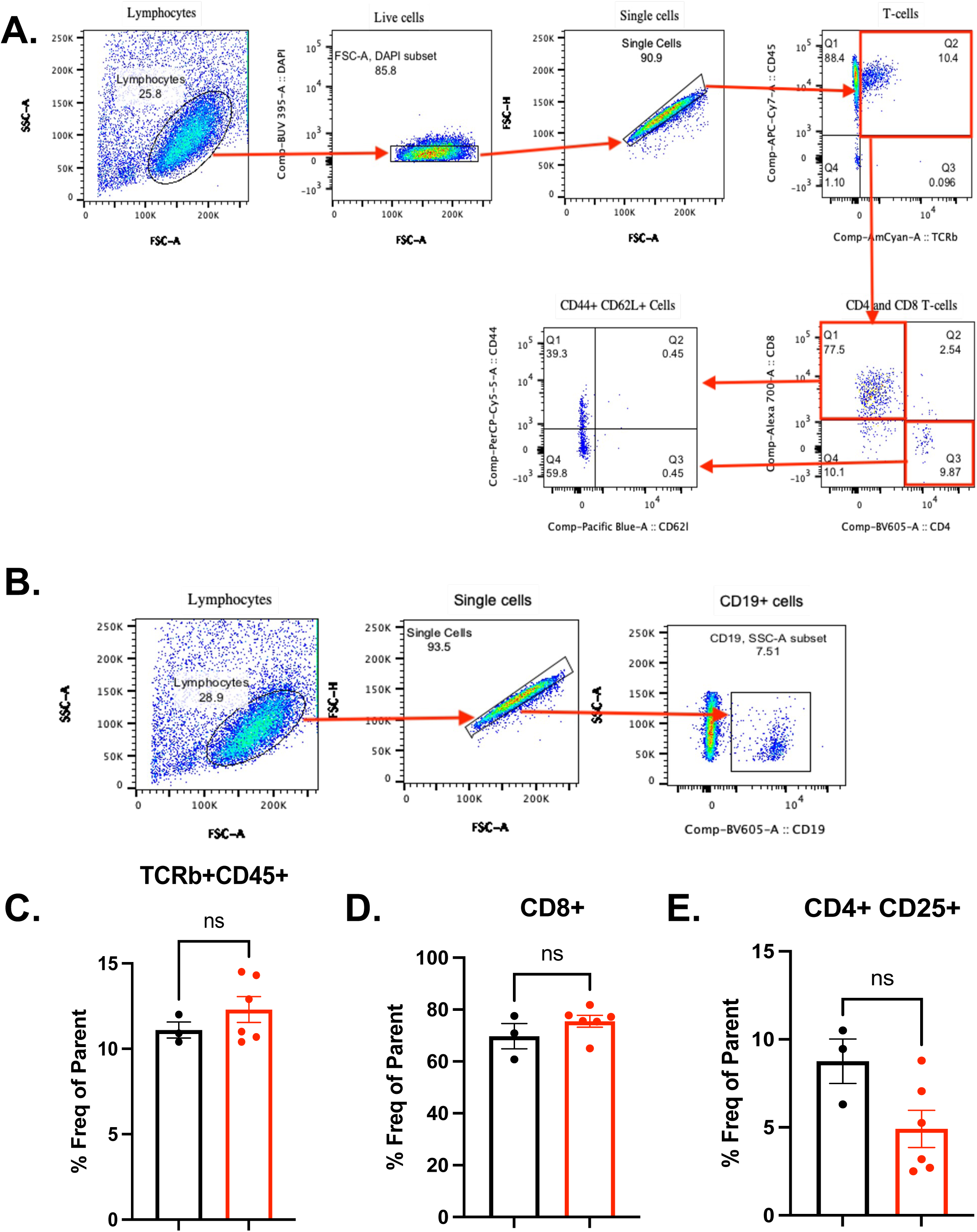
Gating Strategy and T-Cell Population in IEL of female NOD mice. **(A)** Gating strategy for T-cell populations **(B)** B-cell population. **(C)** TCRβ+ CD45+ cell population represents a % fraction of CD45+ cells. **(D)** CD8+ T-cells population represents a % fraction of TCRβ+ CD45+ cells. **(E)** CD4+ CD25+ T-cell population represents as a % fraction of CD4+ population. Statistical analysis was performed using one-way ANOVA for insulitis and two-tailed unpaired Student’s t-test for the gene expression.

**Fig S3:**
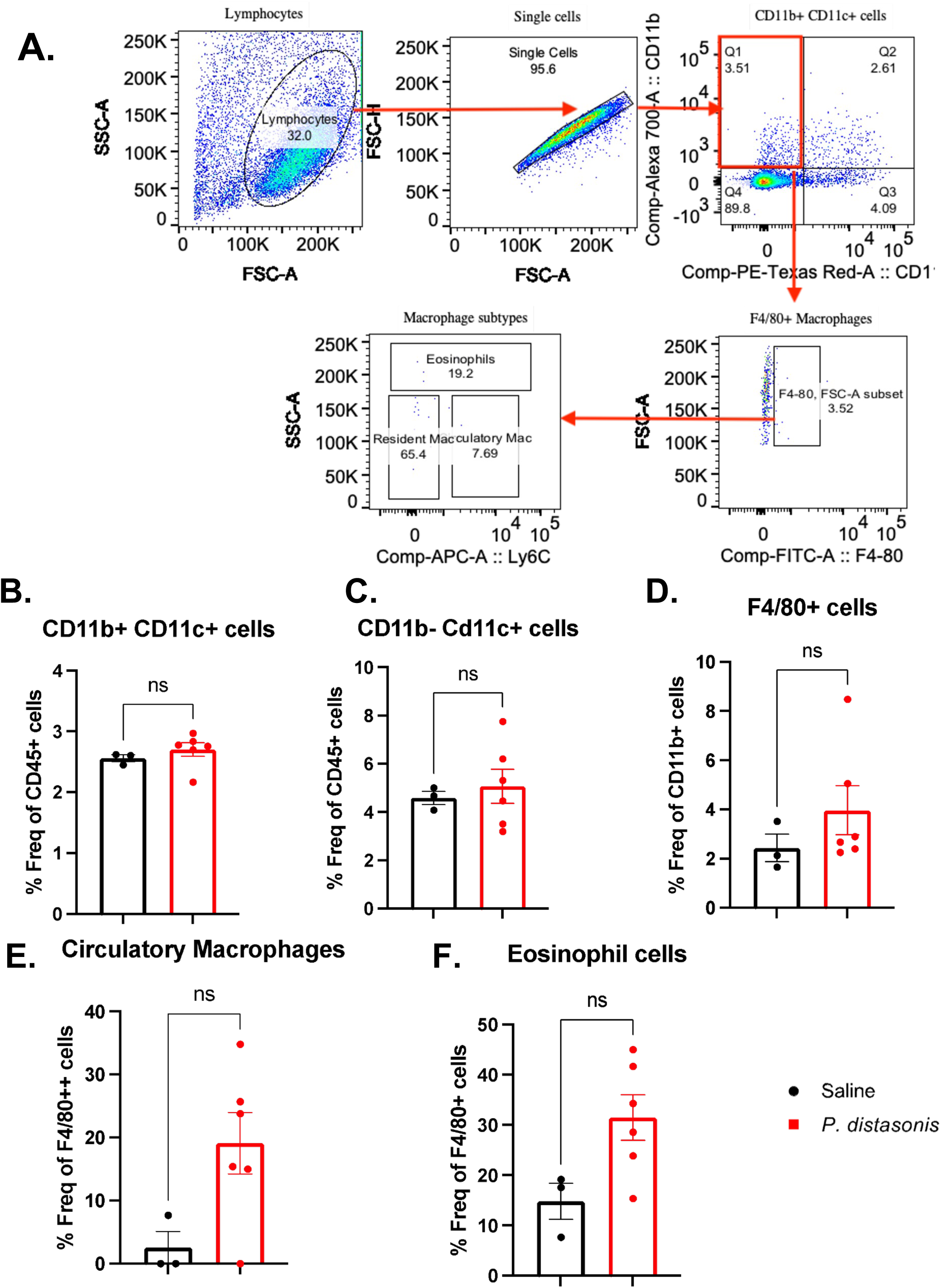
Gating Strategy and Innate Cell Population in IEL of female NOD mice. **(A)** Gating Strategy for innate immune-cell population. **(B)** CD11b+ CD11c+ Dendritic cells, **(C)** CD11b-CD11c+ Dendritic cells in % of CD45+ cells. **(D)** Total F4/80+ macrophages in CD11b+ cell population. **(E)** Circulatory Macrophages **(D)** Eosinophils represent a % fraction of F4/80+ cells. Statistical analysis was performed using one-way ANOVA for insulitis and two-tailed unpaired Student’s t-test for the gene expression.

**Fig S4:**
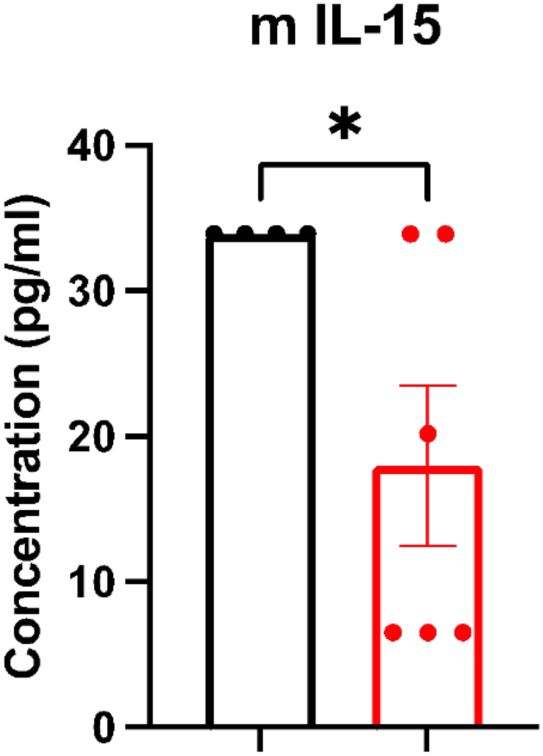
Cytokine profile of *P. distasonis* colonized mice compared to control: IL-15 Cytokine concentration in 12-week *P. distasonis* colonized NOD female mice. saline (n=8), *P. distasonis* (n=9). Statistical analysis was performed using one-way ANOVA for insulitis and two-tailed unpaired Student’s t-test for the gene expression.

## REFERENCES

1. Pugliese A. Autoreactive T cells in type 1 diabetes. J Clin Invest 2017;127(8):2881–91. doi: 10.1172/JCI94549 [published Online First: 2017/08/02]

2. Patterson CC, Harjutsalo V, Rosenbauer J, et al. Trends and cyclical variation in the incidence of childhood type 1 diabetes in 26 European centres in the 25 year period 1989-2013: a multicentre prospective registration study. Diabetologia 2019;62(3):408–17. doi: 10.1007/s00125-018-4763-3 [published Online First: 2018/11/30]

3. Mayer-Davis EJ, Lawrence JM, Dabelea D, et al. Incidence Trends of Type 1 and Type 2 Diabetes among Youths, 2002-2012. N Engl J Med 2017;376(15):1419–29. doi: 10.1056/NEJMoa1610187 [published Online First: 2017/04/14]

4. Pociot F. Type 1 diabetes genome-wide association studies: not to be lost in translation. Clin Transl Immunology 2017;6(12):e162. doi: 10.1038/cti.2017.51 [published Online First: 2018/01/16]

5. Op de Beeck A, Eizirik DL. Viral infections in type 1 diabetes mellitus--why the beta cells? Nat Rev Endocrinol 2016;12(5):263–73. doi: 10.1038/nrendo.2016.30

6. Christen U, Hintermann E, Holdener M, et al. Viral triggers for autoimmunity: is the ‘glass of molecular mimicry’ half full or half empty? J Autoimmun 2010;34(1):38–44. doi: 10.1016/j.jaut.2009.08.001

7. Rewers M, Ludvigsson J. Environmental risk factors for type 1 diabetes. Lancet 2016;387(10035):2340–48. doi: 10.1016/S0140-6736(16)30507-4 [published Online First: 2016/06/16]

8. Rewers M, Hyoty H, Lernmark A, et al. The Environmental Determinants of Diabetes in the Young (TEDDY) Study: 2018 Update. Curr Diab Rep 2018;18(12):136. doi: 10.1007/s11892-018-1113-2 [published Online First: 2018/10/26]

9. Maynard CL, Elson CO, Hatton RD, et al. Reciprocal interactions of the intestinal microbiota and immune system. Nature 2012;489(7415):231–41. doi: 10.1038/nature11551 [published Online First: 2012/09/14]

10. Korpela K, Helve O, Kolho KL, et al. Maternal Fecal Microbiota Transplantation in Cesarean-Born Infants Rapidly Restores Normal Gut Microbial Development: A Proof-of-Concept Study. Cell 2020;183(2):324–34 e5. doi: 10.1016/j.cell.2020.08.047 [published Online First: 2020/10/03]

11. Dedrick S, Sundaresh B, Huang Q, et al. The Role of Gut Microbiota and Environmental Factors in Type 1 Diabetes Pathogenesis. Front Endocrinol (Lausanne) 2020;11:78. doi: 10.3389/fendo.2020.00078 [published Online First: 2020/03/17]

12. Kostic AD, Gevers D, Siljander H, et al. The dynamics of the human infant gut microbiome in development and in progression toward type 1 diabetes. Cell Host Microbe 2015;17(2):260–73. doi: 10.1016/j.chom.2015.01.001 [published Online First: 2015/02/11]

13. Vatanen T, Kostic AD, d’Hennezel E, et al. Variation in Microbiome LPS Immunogenicity Contributes to Autoimmunity in Humans. Cell 2016;165(4):842–53. doi: 10.1016/j.cell.2016.04.007 [published Online First: 2016/05/03]

14. Vatanen T, Plichta DR, Somani J, et al. Genomic variation and strain-specific functional adaptation in the human gut microbiome during early life. Nat Microbiol 2019;4(3):470–79. doi: 10.1038/s41564-018-0321-5 [published Online First: 2018/12/19]

15. Stewart CJ, Ajami NJ, O’Brien JL, et al. Temporal development of the gut microbiome in early childhood from the TEDDY study. Nature 2018;562(7728):583–88. doi: 10.1038/s41586-018-0617-x [published Online First: 2018/10/26]

16. Vatanen T, de Beaufort C, Marcovecchio ML, et al. Gut microbiome shifts in people with type 1 diabetes are associated with glycaemic control: an INNODIA study. Diabetologia 2024 doi: 10.1007/s00125-024-06192-7 [published Online First: 2024/06/04]

17. Sun H, Guo Y, Wang H, et al. Gut commensal Parabacteroides distasonis alleviates inflammatory arthritis. Gut 2023;72(9):1664–77. doi: 10.1136/gutjnl-2022-327756 [published Online First: 2023/01/06]

18. Cuffaro B, Assohoun ALW, Boutillier D, et al. In Vitro Characterization of Gut Microbiota-Derived Commensal Strains: Selection of Parabacteroides distasonis Strains Alleviating TNBS-Induced Colitis in Mice. Cells 2020;9(9) doi: 10.3390/cells9092104 [published Online First: 2020/09/20]

19. Liu D, Zhang S, Li S, et al. Indoleacrylic acid produced by Parabacteroides distasonis alleviates type 2 diabetes via activation of AhR to repair intestinal barrier. BMC Biol 2023;21(1):90. doi: 10.1186/s12915-023-01578-2 [published Online First: 2023/04/19]

20. Cuffaro B, Boutillier D, Desramaut J, et al. Characterization of Two Parabacteroides distasonis Candidate Strains as New Live Biotherapeutics against Obesity. Cells 2023;12(9) doi: 10.3390/cells12091260 [published Online First: 2023/05/13]

21. Wei W, Wong CC, Jia Z, et al. Parabacteroides distasonis uses dietary inulin to suppress NASH via its metabolite pentadecanoic acid. Nat Microbiol 2023;8(8):1534–48. doi: 10.1038/s41564-023-01418-7 [published Online First: 2023/06/30]

22. Gervason S, Meleine M, Lolignier S, et al. Antihyperalgesic properties of gut microbiota: Parabacteroides distasonis as a new probiotic strategy to alleviate chronic abdominal pain. Pain 2024;165(5):e39–e54. doi: 10.1097/j.pain.0000000000003075 [published Online First: 2023/09/27]

23. Koh GY, Kane A, Lee K, et al. Parabacteroides distasonis attenuates toll-like receptor 4 signaling and Akt activation and blocks colon tumor formation in high-fat diet-fed azoxymethane-treated mice. Int J Cancer 2018;143(7):1797–805. doi: 10.1002/ijc.31559 [published Online First: 2018/04/27]

24. benlin Wang YQ, Ming Xie, Pengcheng Huang, Yao Yu, Qi Sun, Wentai Shangguan, Weijia Li, Zhangrui Zhu, Jingwen Xue, Zhengyuan Feng, Yuexuan Zhu, Qishen Yang, Peng Wu. Gut microbiota Parabacteroides distasonis enchances the efficacy of immunotherapy for bladder cancer by activating anti-tumor immune responses, 2023.

25. Koh GY, Kane AV, Wu X, et al. Parabacteroides distasonis attenuates tumorigenesis, modulates inflammatory markers and promotes intestinal barrier integrity in azoxymethane-treated A/J mice. Carcinogenesis 2020;41(7):909–17. doi: 10.1093/carcin/bgaa018 [published Online First: 2020/03/03]

26. Girdhar K, Huang Q, Chow IT, et al. A gut microbial peptide and molecular mimicry in the pathogenesis of type 1 diabetes. Proc Natl Acad Sci U S A 2022;119(31):e2120028119. doi: 10.1073/pnas.2120028119 [published Online First: 2022/07/26]

27. Daniel D, Gill RG, Schloot N, et al. Epitope specificity, cytokine production profile and diabetogenic activity of insulin-specific T cell clones isolated from NOD mice. Eur J Immunol 1995;25(4):1056–62. doi: 10.1002/eji.1830250430

28. Pathiraja V, Kuehlich JP, Campbell PD, et al. Proinsulin-specific, HLA-DQ8, and HLA-DQ8-transdimer-restricted CD4+ T cells infiltrate islets in type 1 diabetes. Diabetes 2015;64(1):172–82. doi: 10.2337/db14-0858

29. Michels AW, Landry LG, McDaniel KA, et al. Islet-Derived CD4 T Cells Targeting Proinsulin in Human Autoimmune Diabetes. Diabetes 2017;66(3):722–34. doi: 10.2337/db16-1025 [published Online First: 2016/12/07]

30. Alleva DG, Crowe PD, Jin L, et al. A disease-associated cellular immune response in type 1 diabetics to an immunodominant epitope of insulin. J Clin Invest 2001;107(2):173–80. doi: 10.1172/JCI8525

31. Nakayama M, McDaniel K, Fitzgerald-Miller L, et al. Regulatory vs. inflammatory cytokine T-cell responses to mutated insulin peptides in healthy and type 1 diabetic subjects. Proc Natl Acad Sci U S A 2015;112(14):4429–34. doi: 10.1073/pnas.1502967112 [published Online First: 2015/04/02]

32. Yang J, Chow IT, Sosinowski T, et al. Autoreactive T cells specific for insulin B:11-23 recognize a low-affinity peptide register in human subjects with autoimmune diabetes. Proc Natl Acad Sci U S A 2014;111(41):14840–5. doi: 10.1073/pnas.1416864111

33. Unanue ER. Antigen presentation in the autoimmune diabetes of the NOD mouse. Annu Rev Immunol 2014;32:579–608. doi: 10.1146/annurev-immunol-032712-095941 [published Online First: 2014/02/07]

34. Monsted MO, Falck ND, Pedersen K, et al. Intestinal permeability in type 1 diabetes: An updated comprehensive overview. J Autoimmun 2021;122:102674. doi: 10.1016/j.jaut.2021.102674 [published Online First: 2021/06/29]

35. Bell KJ, Saad S, Tillett BJ, et al. Metabolite-based dietary supplementation in human type 1 diabetes is associated with microbiota and immune modulation. Microbiome 2022;10(1):9. doi: 10.1186/s40168-021-01193-9 [published Online First: 2022/01/21]

36. Schwarzer M, Makki K, Storelli G, et al. Lactobacillus plantarum strain maintains growth of infant mice during chronic undernutrition. Science 2016;351(6275):854–7. doi: 10.1126/science.aad8588 [published Online First: 2016/02/26]

37. Girdhar K, Dogru YD, Huang Q, et al. Dynamics of the gut microbiome, IgA response, and plasma metabolome in the development of pediatric celiac disease. Microbiome 2023;11(1):9. doi: 10.1186/s40168-022-01429-2 [published Online First: 2023/01/14]

38. Liu CM, Aziz M, Kachur S, et al. BactQuant: an enhanced broad-coverage bacterial quantitative real-time PCR assay. BMC Microbiol 2012;12:56. doi: 10.1186/1471-2180-12-56 [published Online First: 2012/04/19]

39. Kozich JJ, Westcott SL, Baxter NT, et al. Development of a dual-index sequencing strategy and curation pipeline for analyzing amplicon sequence data on the MiSeq Illumina sequencing platform. Appl Environ Microbiol 2013;79(17):5112–20. doi: 10.1128/AEM.01043-13 [published Online First: 2013/06/25]

40. Callahan BJ, McMurdie PJ, Rosen MJ, et al. DADA2: High-resolution sample inference from Illumina amplicon data. Nat Methods 2016;13(7):581–3. doi: 10.1038/nmeth.3869 [published Online First: 2016/05/24]

41. Cole JR, Wang Q, Fish JA, et al. Ribosomal Database Project: data and tools for high throughput rRNA analysis. Nucleic Acids Res 2014;42(Database issue):D633–42. doi: 10.1093/nar/gkt1244 [published Online First: 2013/11/30]

42. McMurdie PJ, Holmes S. phyloseq: an R package for reproducible interactive analysis and graphics of microbiome census data. PLoS One 2013;8(4):e61217. doi: 10.1371/journal.pone.0061217 [published Online First: 2013/05/01]

43. Robinson MD, McCarthy DJ, Smyth GK. edgeR: a Bioconductor package for differential expression analysis of digital gene expression data. Bioinformatics 2010;26(1):139–40. doi: 10.1093/bioinformatics/btp616 [published Online First: 2009/11/17]

44. Glickman ME, Rao SR, Schultz MR. False discovery rate control is a recommended alternative to Bonferroni-type adjustments in health studies. J Clin Epidemiol 2014;67(8):850–7. doi: 10.1016/j.jclinepi.2014.03.012 [published Online First: 2014/05/17]

45. Liu Y, Dai M. Trimethylamine N-Oxide Generated by the Gut Microbiota Is Associated with Vascular Inflammation: New Insights into Atherosclerosis. Mediators Inflamm 2020;2020:4634172. doi: 10.1155/2020/4634172 [published Online First: 2020/03/10]

46. Seldin MM, Meng Y, Qi H, et al. Trimethylamine N-Oxide Promotes Vascular Inflammation Through Signaling of Mitogen-Activated Protein Kinase and Nuclear Factor-kappaB. J Am Heart Assoc 2016;5(2) doi: 10.1161/JAHA.115.002767 [published Online First: 2016/02/24]

47. Zhang Y, Wang Y, Ke B, et al. TMAO: how gut microbiota contributes to heart failure. Transl Res 2021;228:109–25. doi: 10.1016/j.trsl.2020.08.007 [published Online First: 2020/08/26]

48. Dolkar P, Deyang T, Anand N, et al. Trimethylamine-N-oxide and cerebral stroke risk: A review. Neurobiol Dis 2024;192:106423. doi: 10.1016/j.nbd.2024.106423 [published Online First: 2024/01/30]

49. Saaoud F, Liu L, Xu K, et al. Aorta- and liver-generated TMAO enhances trained immunity for increased inflammation via ER stress/mitochondrial ROS/glycolysis pathways. JCI Insight 2023;8(1) doi: 10.1172/jci.insight.158183 [published Online First: 2022/11/18]

50. Arffman M, Hakkarainen P, Keskimaki I, et al. Long-term and recent trends in survival and life expectancy for people with type 1 diabetes in Finland. Diabetes Res Clin Pract 2023;198:110580. doi: 10.1016/j.diabres.2023.110580 [published Online First: 2023/02/22]

51. Abraham C, Cho JH. Inflammatory bowel disease. N Engl J Med 2009;361(21):2066–78. doi: 10.1056/NEJMra0804647 [published Online First: 2009/11/20]

52. Shale M, Schiering C, Powrie F. CD4(+) T-cell subsets in intestinal inflammation. Immunol Rev 2013;252(1):164–82. doi: 10.1111/imr.12039 [published Online First: 2013/02/15]

53. Brucklacher-Waldert V, Carr EJ, Linterman MA, et al. Cellular Plasticity of CD4+ T Cells in the Intestine. Front Immunol 2014;5:488. doi: 10.3389/fimmu.2014.00488 [published Online First: 2014/10/24]

54. Lejeune T, Meyer C, Abadie V. B Lymphocytes Contribute to Celiac Disease Pathogenesis. Gastroenterology 2021;160(7):2608–10 e4. doi: 10.1053/j.gastro.2021.02.063 [published Online First: 2021/03/06]

55. Gaifem J, Mendes-Frias A, Wolter M, et al. Akkermansia muciniphila and Parabacteroides distasonis synergistically protect from colitis by promoting ILC3 in the gut. mBio 2024;15(4):e0007824. doi: 10.1128/mbio.00078-24 [published Online First: 2024/03/12]

56. Kverka M, Zakostelska Z, Klimesova K, et al. Oral administration of Parabacteroides distasonis antigens attenuates experimental murine colitis through modulation of immunity and microbiota composition. Clin Exp Immunol 2011;163(2):250–9. doi: 10.1111/j.1365-2249.2010.04286.x [published Online First: 2010/11/23]

57. Fletcher CM, Coyne MJ, Bentley DL, et al. Phase-variable expression of a family of glycoproteins imparts a dynamic surface to a symbiont in its human intestinal ecosystem. Proc Natl Acad Sci U S A 2007;104(7):2413–8. doi: 10.1073/pnas.0608797104 [published Online First: 2007/02/08]

58. Wang K, Liao M, Zhou N, et al. Parabacteroides distasonis Alleviates Obesity and Metabolic Dysfunctions via Production of Succinate and Secondary Bile Acids. Cell Rep 2019;26(1):222–35 e5. doi: 10.1016/j.celrep.2018.12.028 [published Online First: 2019/01/04]

59. Lopetuso LR, Petito V, Graziani C, et al. Gut Microbiota in Health, Diverticular Disease, Irritable Bowel Syndrome, and Inflammatory Bowel Diseases: Time for Microbial Marker of Gastrointestinal Disorders. Dig Dis 2018;36(1):56–65. doi: 10.1159/000477205 [published Online First: 2017/07/07]

60. Okayasu I, Hatakeyama S, Yamada M, et al. A novel method in the induction of reliable experimental acute and chronic ulcerative colitis in mice. Gastroenterology 1990;98(3):694–702. doi: 10.1016/0016-5085(90)90290-h [published Online First: 1990/03/01]

61. Ezeji JC, Sarikonda DK, Hopperton A, et al. Parabacteroides distasonis: intriguing aerotolerant gut anaerobe with emerging antimicrobial resistance and pathogenic and probiotic roles in human health. Gut Microbes 2021;13(1):1922241. doi: 10.1080/19490976.2021.1922241 [published Online First: 2021/07/02]

62. Gonzalez-Paez GE, Roncase EJ, Wolan DW. X-ray structure of an inactive zymogen clostripain-like protease from Parabacteroides distasonis. Acta Crystallogr D Struct Biol 2019;75(Pt 3):325–32. doi: 10.1107/S2059798319000809 [published Online First: 2019/04/06]

63. Xu JH, Jiang Z, Solania A, et al. A Commensal Dipeptidyl Aminopeptidase with Specificity for N-Terminal Glycine Degrades Human-Produced Antimicrobial Peptides in Vitro. ACS Chem Biol 2018;13(9):2513–21. doi: 10.1021/acschembio.8b00420 [published Online First: 2018/08/08]

64. Mason D. A very high level of crossreactivity is an essential feature of the T-cell receptor. Immunol Today 1998;19(9):395–404. doi: 10.1016/s0167-5699(98)01299-7 [published Online First: 1998/09/24]

65. Calis JJ, de Boer RJ, Kesmir C. Degenerate T-cell recognition of peptides on MHC molecules creates large holes in the T-cell repertoire. PLoS Comput Biol 2012;8(3):e1002412. doi: 10.1371/journal.pcbi.1002412 [published Online First: 2012/03/08]

66. Wucherpfennig KW, Strominger JL. Molecular mimicry in T cell-mediated autoimmunity: viral peptides activate human T cell clones specific for myelin basic protein. Cell 1995;80(5):695–705. doi: 10.1016/0092-8674(95)90348-8 [published Online First: 1995/03/10]

67. Mana P, Goodyear M, Bernard C, et al. Tolerance induction by molecular mimicry: prevention and suppression of experimental autoimmune encephalomyelitis with the milk protein butyrophilin. Int Immunol 2004;16(3):489–99. doi: 10.1093/intimm/dxh049 [published Online First: 2004/02/24]

68. Greiling TM, Dehner C, Chen X, et al. Commensal orthologs of the human autoantigen Ro60 as triggers of autoimmunity in lupus. Sci Transl Med 2018;10(434) doi: 10.1126/scitranslmed.aan2306 [published Online First: 2018/03/30]

69. Gil-Cruz C, Perez-Shibayama C, De Martin A, et al. Microbiota-derived peptide mimics drive lethal inflammatory cardiomyopathy. Science 2019;366(6467):881–86. doi: 10.1126/science.aav3487 [published Online First: 2019/11/16]

70. Harkiolaki M, Holmes SL, Svendsen P, et al. T cell-mediated autoimmune disease due to low-affinity crossreactivity to common microbial peptides. Immunity 2009;30(3):348–57. doi: 10.1016/j.immuni.2009.01.009 [published Online First: 2009/03/24]

71. Tian J, Lehmann PV, Kaufman DL. T cell cross-reactivity between coxsackievirus and glutamate decarboxylase is associated with a murine diabetes susceptibility allele. J Exp Med 1994;180(5):1979–84. doi: 10.1084/jem.180.5.1979 [published Online First: 1994/11/01]

72. Atkinson MA, Bowman MA, Campbell L, et al. Cellular immunity to a determinant common to glutamate decarboxylase and coxsackie virus in insulin-dependent diabetes. J Clin Invest 1994;94(5):2125–9. doi: 10.1172/JCI117567 [published Online First: 1994/11/01]

73. Tai N, Peng J, Liu F, et al. Microbial antigen mimics activate diabetogenic CD8 T cells in NOD mice. J Exp Med 2016;213(10):2129–46. doi: 10.1084/jem.20160526 [published Online First: 2016/09/14]

74. Honeyman MC, Stone NL, Falk BA, et al. Evidence for molecular mimicry between human T cell epitopes in rotavirus and pancreatic islet autoantigens. J Immunol 2010;184(4):2204–10. doi: 10.4049/jimmunol.0900709 [published Online First: 2010/01/20]

75. Huang Q, Chow I-T, Brady C, et al. Parabacteroides distasonis insulin B:9-23 epitope mimic stimulates insulin specific T-cells and enhances Type 1 Diabetes in NOD mice. bioRxiv 2020:2020.10.22.350801. doi: 10.1101/2020.10.22.350801

76. Lanz TV, Brewer RC, Ho PP, et al. Clonally expanded B cells in multiple sclerosis bind EBV EBNA1 and GlialCAM. Nature 2022;603(7900):321–27. doi: 10.1038/s41586-022-04432-7 [published Online First: 2022/01/25]

77. Bjornevik K, Cortese M, Healy BC, et al. Longitudinal analysis reveals high prevalence of Epstein-Barr virus associated with multiple sclerosis. Science 2022;375(6578):296–301. doi: 10.1126/science.abj8222 [published Online First: 2022/01/14]

78. Kent SC, Mannering SI, Michels AW, et al. Deciphering the Pathogenesis of Human Type 1 Diabetes (T1D) by Interrogating T Cells from the “Scene of the Crime“. Curr Diab Rep 2017;17(10):95. doi: 10.1007/s11892-017-0915-y [published Online First: 2017/09/03]

79. James EA, Mallone R, Kent SC, et al. T-Cell Epitopes and Neo-epitopes in Type 1 Diabetes: A Comprehensive Update and Reappraisal. Diabetes 2020;69(7):1311–35. doi: 10.2337/dbi19-0022 [published Online First: 2020/06/21]

